# The SARS-CoV-2 transcriptome and the dynamics of the S gene furin cleavage site in primary human airway epithelia

**DOI:** 10.1101/2021.02.03.429670

**Authors:** Wei Zou, Min Xiong, Siyuan Hao, Elizabeth Yan Zhang-Chen, Nathalie Baumlin, Michael D. Kim, Matthias Salathe, Ziying Yan, Jianming Qiu

## Abstract

The novel severe acute respiratory syndrome coronavirus-2 (SARS-CoV-2) caused the devastating ongoing coronavirus disease-2019 (COVID-19) pandemic which poses a great threat to global public health. The spike (S) polypeptide of SARS-CoV-2 consists of the S1 and S2 subunits and is processed by cellular proteases at the S1/S2 boundary. The inclusion of the 4 amino acids (PRRA) at the S1/S2 boundary forms a furin cleavage site (FCS), ^682^RRAR↓S^686^, distinguishing SARS-CoV-2 from its closest relative, the SARS-CoV. Various deletions surrounding the FCS have been identified in patients. When SARS-CoV-2 propagated in Vero cells, the virus acquired various deletions surrounding the FCS. In the present study, we studied the viral transcriptome in SARS-CoV-2 infected primary human airway epithelia (HAE) cultured at an air-liquid interface (ALI) with an emphasis on the viral genome stability at the S1/S2 boundary using RNA-seq. While we found overall the viral transcriptome is similar to that generated from infected Vero cells, we identified a high percentage of mutated viral genome and transcripts in HAE-ALI. Two highly frequent deletions were found at the S1/S2 boundary of the S gene: one is a deletion of 12 amino acids, ^678^TNSP<u>RRAR↓S</u>VAS^689^, which contains the FCS, another is a deletion of 5 amino acids, ^675^QTQTN^679^, which is two amino acids upstream of the FCS. Further studies on the dynamics of the FCS deletions in apically released virions revealed that the selective pressure for the FCS maintains the S gene stability in HAE-ALI but with exceptions, in which the FCS deletions are remained at a high rate. Thus, our study presents evidence for the role of unique properties of human airway epithelia in the dynamics of the FCS region during infection of human airways, which is donor-dependent.

## Introduction

The ongoing coronavirus disease-2019 (COVID-19) outbreak, caused by the novel severe acute respiratory syndrome coronavirus-2 (SARS-CoV-2), poses a great threat to global public health with a devastating mortality (1-4). The virus has spread with unprecedented speed and has infected >100 million people worldwide, causing so far >2 million deaths. The efficacy of the only United States Food and Drug Administration (FDA) approved drug, VEKLURY (remdesivir), to treat COVID-19 patients is limited to early phases of the disease and is supportive (5). Ongoing rollout of the two mRNA-based COVID-19 vaccines approved by the FDA under an emergency use authorization, with more vaccines to follow soon, is the hope to prevent COVID-19 and contain the virus (6).

SARS-CoV-2 phylogenetically belongs to the genus *Betacoronavirus* of the family *Coronaviridae* (7,8), and is closely related to the previously identified severe acute respiratory syndrome coronavirus (SARS-CoV) with an identity of 79% in genome sequence (9,10). SARS had an outbreak in 2002-2003 (11,12). The genome organization of SARS-CoV-2 is the same as other betacoronaviruses. It has six major open reading frames (ORFs) arranged in order from the structured 5’ untranslated region (UTR) to 3’ UTR (13,14): replicases (ORF1a and ORF1b), spike (S), envelope (E), membrane (M), and nucleocapsid (N). In addition, at least seven ORFs encoding accessory proteins (3a, 6, 7a, 7b, 8, 9a, and 9b) are interspersed between the structural protein genes (10,15).

The replication and transcription of SARS-CoV-2 largely resemble that of the SARS-CoV (15,16). The accepted model for coronavirus transcription indicates that all viral mRNAs have a common 5’-leader (L) sequence or the 5’-cap structure at the 5’UTR, and a common poly(A) tail at the 3UTR (15,17-19). The highly conserved leader sequences contain the transcription regulatory sequences (TRS) which play an important role in viral RNA transcription (18-20). Upon cell entry of the virus, the incoming positive-sense genomic RNA (+gRNA) subjects it to immediate translation of two large ORFs, ORF1a and ORF1b, for viral nonstructural proteins, which form the viral replication and transcription complex (RTC) (21). In the complex, the viral +gRNA serves as the template for the production of negative-sense (-)gRNA and sub-genome RNA (-sgRNA) intermediates, which in turn serve as the templates for the synthesis of +gRNA and +sgRNAs (22). The positive-sense viral RNAs are the mRNAs used for the translations of at least 20 viral nonstructural proteins and 5 structural proteins, spike protein S, envelope protein E, membrane protein M, and nucleocapsid protein N (21). Next, the newly synthesized +gRNA is encapsidated by the N protein to assemble progeny virions with other viral structural proteins, M, E, and S (21).

Most SARS-CoV-2 structural and nonstructural proteins share greater than 85% identity in protein sequence with SARS-CoV, whereas their S proteins only share an identity of approximately 77% (2). S protein consists of two subunits, S1 and S2, and is a key glycoprotein responsible for receptor binding and determining the host tropism, pathogenicity, and transmissibility (23,24). It forms a homotrimer on the virion surface and triggers viral entry into target cells via binding of the S1 subunit to its cognate receptor, angiotensin-converting enzyme 2 (ACE2) (2,25,26). One significant difference among S proteins of SARS-CoV-2, SARS-CoV, and other bat SARS-like coronaviruses, such as BtCoV-RaTG13, is the addition of 4 amino acids, PRRA, at the S1/S2 boundary (24). This insertion forms a polybasic residue motif, assembling a furin cleavage site (FCS), RRAR↓S, which is highly related to the furin cleavage consensus sequence RX[K/R]R (X, any amino acid) (27). The absence of the FCS in the other betacoronaviruses suggests the insertion of PRRA is a key factor in the virulence of SARS-CoV-2, which has been shown to broaden cell tropism, transmissibility, and pathogenicity of the virus (28-30).

Viral transcriptomes of SARS-CoV-2 have been studied by several groups but only in infected Vero cells (15,17), which revealed quick mutations in the S1/S2 boundary of the S gene, including the loss of the FCS and the immediately adjacent amino acids upstream or downstream of the FCS. The loss of the FCS has been identified in progeny virions replicated in Vero cells (17,31-34). The mutant viruses were stable, quickly took over the wild-type (WT) virus, and became the dominant population during passaging. Of note, various deletions surrounding the FCS have been identified in patients. This raises the question of how the FCS region deletions are selected in human airways.

In this study, we used RNA-seq to analyze the viral transcriptome of SARS-CoV-2 in the infected human airway epithelia (HAE) cultured at an air-liquid interface (HAE-ALI), which mimics natural viral infection of human airways (35,36). While the viral transcriptome overall recapitulated that in Vero cells, we discovered that there is a selective pressure in HAE-ALI to suppress the deletions at the S1/S2 boundary and that this pressure appears individual donor dependent. We identified two FCS region deletions that are strikingly amplified in two HAE-ALI cultures after 2-3 weeks of infection, whereas these deletions were suppressed in five other HAE-ALI cultures.

## Materials and Methods

### Viruses

SARS-CoV-2 (NR-52281), isolate USA-WA1/2020 (Batch no.: 70034262), was obtained from BEI Resources (Manassas, VA) and designated as P0 passage. The virus used for infections of HAE-ALI was propagated once in Vero-E6 cells, designated as P1 passage. Viruses were titrated by plaque assays on Vero-E6 cells and stored at −80°C as previously described (36). A biosafety protocol to work on SARS-CoV-2 infection in the biosafety level-3 (BSL3) lab was approved by the Institutional Biosafety Committee of the University of Kansas Medical Center.

### HAE-ALI cultures

Primary HAE-ALI cultures, lots of B2-20, B3-20, B4-20, B9-20, B15-20, and B16-20, were prepared from bronchial airway epithelial cells isolated from various donors. They were obtained from the Cells and Tissue Core of the Center for Gene Therapy, University of Iowa and polarized in Transwell inserts (0.33 cm^2^; Costar, Corning, Tewksbury, WA). L209 and KC19 HAE-ALI cultures were prepared from propagated bronchial airway cells of the L209 and KC19 donors provided by the Department of Internal Medicine, University of Kansas Medical Center. They were polarized on Transwell inserts (1.1 cm^2^; Costar, Corning). The HAE-ALI cultures that had transepithelial electrical resistance (TEER) of > 1,000 Ω·cm^2^, determined with an epithelial volt-ohm meter (MilliporeSigma, Burlington, MA), were used for infections.

### Virus infections

Polarized HAE-ALI cultures were infected with SARS-CoV-2 at a multiplicity of infection (MOI) of 0.2 or 2. The inoculum of 100 µl or 300 µl was apically applied to the 0.33 cm^2^ or 1.1 cm^2^ Transwell inserts with an incubation period of 1 h at 37°C and 5% CO_2_. After aspiration of the inoculum, the apical surface of the insert was washed with 100 µl (or 300 µl) of Dulbecco’s phosphate-buffered saline (D-PBS; Corning, Tewksbury, WA) three times to maximally remove the unbound viruses. The HAE-ALI cultures were then placed back into the incubator at 37°C and 5% CO_2_. To collect the apically released progeny from infected cultures, 100 µl (or 300 µl) of D-PBS was added to the apical chamber for 30 min at 37°C and 5% CO_2_. Thereafter, the apical wash was pipetted carefully from the apical chamber.

### Immunofluorescence assay

The membrane of the infected HAE-ALI was cut out and fixed with 4% paraformaldehyde in PBS at 4°C overnight. The fixed membrane was washed in PBS for 5 min three times and then split into several pieces for whole-mount immunostaining. Following permeabilization with 0.2% Triton X-100 for 15 min at room temperature, the slide was incubated with a rabbit monoclonal anti-SARS-CoV-2 nucleocapsid (NP) (# 40143-R001; SinoBiological US, Wayne, PA) at a dilution of 1:25 in PBS with 2% fetal bovine serum for 1 h at 37°C. After washing, the slide was incubated with a rhodamine-conjugated secondary antibody, followed by staining of the nuclei with DAPI (4’,6-diamidino-2-phenylindole).

### RNA extraction

For total RNA extraction, 4 Transwell inserts of HAE-ALI cultures were dissolved in 1 ml of TRIzol Reagent (ThermoFisher, Waltham, MA), following manufacturer’s instructions. Viral RNA was isolated from the virions in apical washes. 50 µl of apical wash was used for the extraction of nuclease digestion-resistant viral RNA using the Quick-RNA Viral kit (#R1035; Zymo Research, Irvine, CA), as described previously (36). The final RNA samples were dissolved in 50 µl of deionized H_2_O and quantified for concentrations using a microplate reader (Synergy H, BioTek).

### RNA-seq

For viral transcriptome, total RNA was extracted from HAE-ALI cultures infected with SARS-CoV-2 at an MOI of 0.2 and 2, respectively, or mock infected at 4 dpi. After RNA quality control and reverse transcription, DNA nanoball sequencing (DNSeq) was performed at BGI Genomics (Cambridge, MA). Briefly, RNA samples were tested using an Agilent 2100 Bioanalyzer (Agilent RNA 6000 Nano Kit). Samples with an RNA Integrity Number (RIN) of ≥ 8.0 were chosen for library construction. rRNA was removed from the total RNA samples by using RNase H or Ribo-Zero method. Then, samples were fragmented in a fragmentation buffer for thermal fragmentation to 130-160 nucleotides (nts). First-strand cDNA was generated by First Strand Mix, then Second Strand Mix was added to synthesize the second-strand cDNA. The reaction product was purified by magnetic beads and end-repaired by addition of adaptors, followed by several rounds of PCR amplification to enrich the cDNA fragments. The PCR products were then purified and subjected to library quality control on the Agilent Technologies 2100 bioanalyzer. The double stranded PCR products were heat denatured and circularized by the splint oligo sequence. The single strand circle DNA (ssCir DNA) were formatted as the final library. The final library was amplified with phi29 to make DNA nanoball (DNB) which have more than 300 copies of one molecule. The DNBs were load into the patterned nanoarray and 2 × 100 paired-end reads were generated in the way of combinatorial Probe-Anchor Synthesis (cPAS).

For RNA-seq of the viral RNAs, the apical washes were collected from infected HAE-ALI cultures at the indicated times (days post-infection, dpi; **Tab. 3**), and were extracted for viral RNA as described above. For library preparation, the stranded-RNA seq kit (Thermo Fisher) was used following the manufacturer’s protocol. The rRNA depletion step was added for the library preparation. The Illumina sequencer NextSeq550 was used to generate pair-end 2 × 150 reads at GeneGoCell Inc. (San Diego, CA).

### PCR amplicon-seq

For sequencing the FCS region of the S gene, viral RNA extracted from the apical washes was reverse-transcribed using AMV (Promega, Madison, WI). A 384-nt sequence covering the S gene FCS region (nt 23,487-23,870) was amplified by PCR of 20 cycles using the primers containing the <u>adaptor</u>sequences: Forward: 5’-<u>ACA CTC TTT CCC TAC ACG ACG CTC TTC CGA TCT</u> TTT TCA AAC ACG TGC AGG C-3’, and Reverse: 5’-<u>GAC TGG AGT TCA GAC GTG TGC TCT TCC GAT CT</u>T CCA GTT AAA GCA CGG TTT AAT-3’. The PCR products were analyzed on 1.5% agarose and excised for purification. The purified DNA samples were quantified on a microplate reader (Synergy LX, BioTek, Winooski, VT), and 500 ng of each DNA sample (20 ng/µl) was sent for PCR amplicon-seq (AMPLICON-EZ) at GENEWIZ, Inc. (South Plainfield, NJ).

### Bioinformatic analyses

#### Total cellular DNBseq data (BGI) and PCR-amplicon-seq (GENEWIZ)

The reads were aligned to the reference SARS-CoV-2 Wuhan-Hu-1 isolate genome (GenBank accession no: MN908947) using BWA v0.7.5a-r405. Sequencing read coverage was calculated using bedtools genomecov of version 2.27.1. We used STAR (2.7.3a) to identify the junction-spanning reads as described previously except that we set the minimal size of deletions as 10 (15).

#### Viral RNA-seq data (GeneGoCell Inc.)

Raw sequence reads (fastq files) were processed through the following steps by the Genenius NGS bioinformatics pipeline (v2.1). Low-quality reads were removed using quality score threshold 25 (Q25). The resulting fastq files were analyzed by FastQC v0.10.1 for quality control (QC). Reads were aligned to the reference genome Wuhan-Hu-1. The alignment results were analyzed using the proprietary GeneGoCell program for variant calling on the target sites as follows: 1) Each read pair was processed to report the variant in the read; 2) Each variant’s allele frequency (AF) was calculated based on # of variant reads / total reads covering the region (both variant and non-variant); 3) Variants with ≥ 1% AF and ≥ 3 variant reads were reported in a variant calling file (vcf). The output of the bioinformatics workflow was collected and further organized/processed in Microsoft Office 365.

#### Data deposition

All the RNA-seq and PCR amplicon-seq data have been deposited in NIH-sponsored BioProject database, PRJNA698337 (https://dataview.ncbi.nlm.nih.gov/object/PRJNA698337?reviewer=k4gtr6eundj03jnpcrj62tq1un).

## Results

### To determine the SARS-CoV-2 transcriptome in SARS-CoV-2 infected HAE-ALI cultures

HAE-ALI^B2-20^ cultures were infected with SARS-CoV-2 at an MOI of 0.2 or 2, or mock infected. At 4 days post-infection (dpi), immunofluorescence assay for the SARS-CoV-2 N protein expression revealed effective SARS-CoV-2 infection in these cultures, with ∼10% and ∼30% of cells positive in the infections at MOIs of 0.2 and 2, respectively (**Fig. 1**). This result was similar to our previous observation (36).

**Fig. 1.**
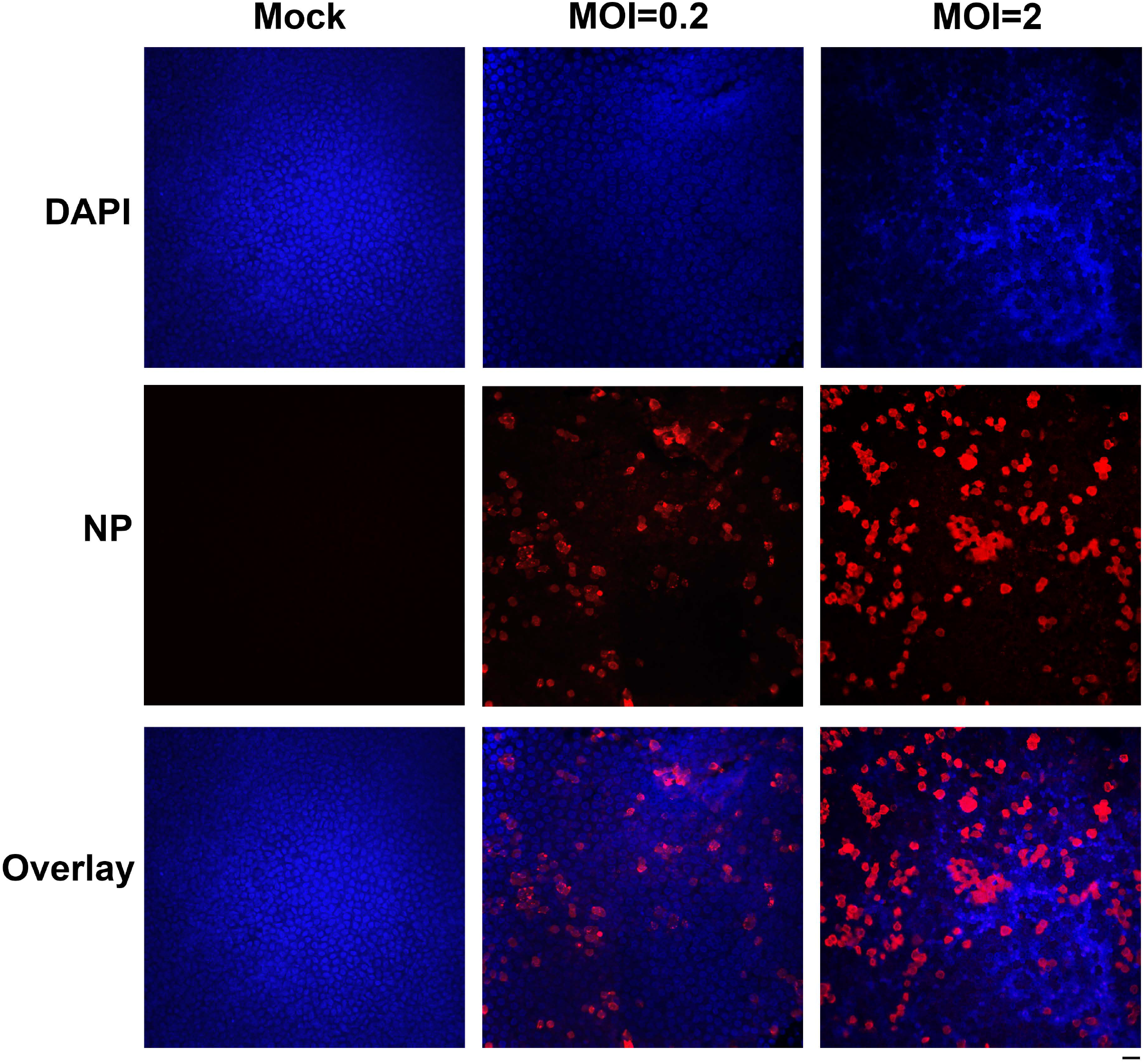
Immunofluorescence analysis of SARS-CoV-2 infection of HAE cells. HAE-ALI^B2-20^ cultures were infected with SARS-CoV-2 at an MOI of 0.2 or 2 pfu/cell, as indicated, mock-infected (Mock). At 4 days post-infection, a piece of the insert membrane was fixed in 4% paraformaldehyde in PBS at 4°C overnight. and subjected to direct immunofluorescence analysis. The membranes were stained with anti-SARS-CoV-2 N protein (NP). Images were taken on a Leica TCS SPE confocal microscope under 40×, which was controlled by Leica Application Suite X software. The nuclei were stained with DAPI (4’=,6-diamidino-2-phenylindole). Scale bar is 20 µM.

Total RNA samples were extracted from infected HAE-ALI cultures at 4 dpi and subjected to reverse transcription, followed by DNA nanoball sequencing. An average total reads of 18.27% and 26.54% were mapped to the SARS-CoV-2 reference genome (Wuhan-Hu-1 isolate; MN908947) in the groups of MOI 0.2 and MOI 2, respectively (**Tab. 1**). No significant difference was observed in the total reads in the two groups. Notably, the RNA-seq data obtained from SARS-CoV-2 infected Vero-E6 cells had up to 70% of the reads mapped to the viral genome (15), which was likely due to the high infectivity of Vero-E6 cells and that not all the cell types in HAE-ALI are permissive to the infection (36). Also, for the whole viral genome coverage, in contrast to the observation in SARS-CoV-2 infected Vero-E6 cells (15), we did not observe an obvious 5’-leader peak in the infected HAE cells (**Fig. 2**). Instead, we observed ∼2-fold higher reads in the 3’-end than that in the 5’-end viral genome.

**Table 1.** Summary of RNA-seq data of SARS-CoV-2 and mock infected HAE-ALI cultures.

| **Samples** | **Total reads** | **Mapped viral reads** | **Mapped viral reads (%)** |
| --- | --- | --- | --- |
| **Mock-1** | 44,690,289 | 0 | 0 |
| **Mock-2** | 44,710,969 | 0 | 0 |
| **Mock-3** | 44,661,039 | 0 | 0 |
| **MOI 0.2-1** | 44,665,846 | 9,109,261 | 20.39 |
| **MOI 0.2-2** | 44,606,709 | 9,446,219 | 21.18 |
| **MOI 0.2-3** | 44,625,565 | 5,907,458 | 13.24 |
| **MOI 2-1** | 44,622,724 | 14,536,455 | 32.58 |
| **MOI 2-2** | 44,658,564 | 12,466,502 | 27.92 |
| **MOI 2-3** | 44,616,157 | 8,529,469 | 19.12 |

**Fig. 2.**
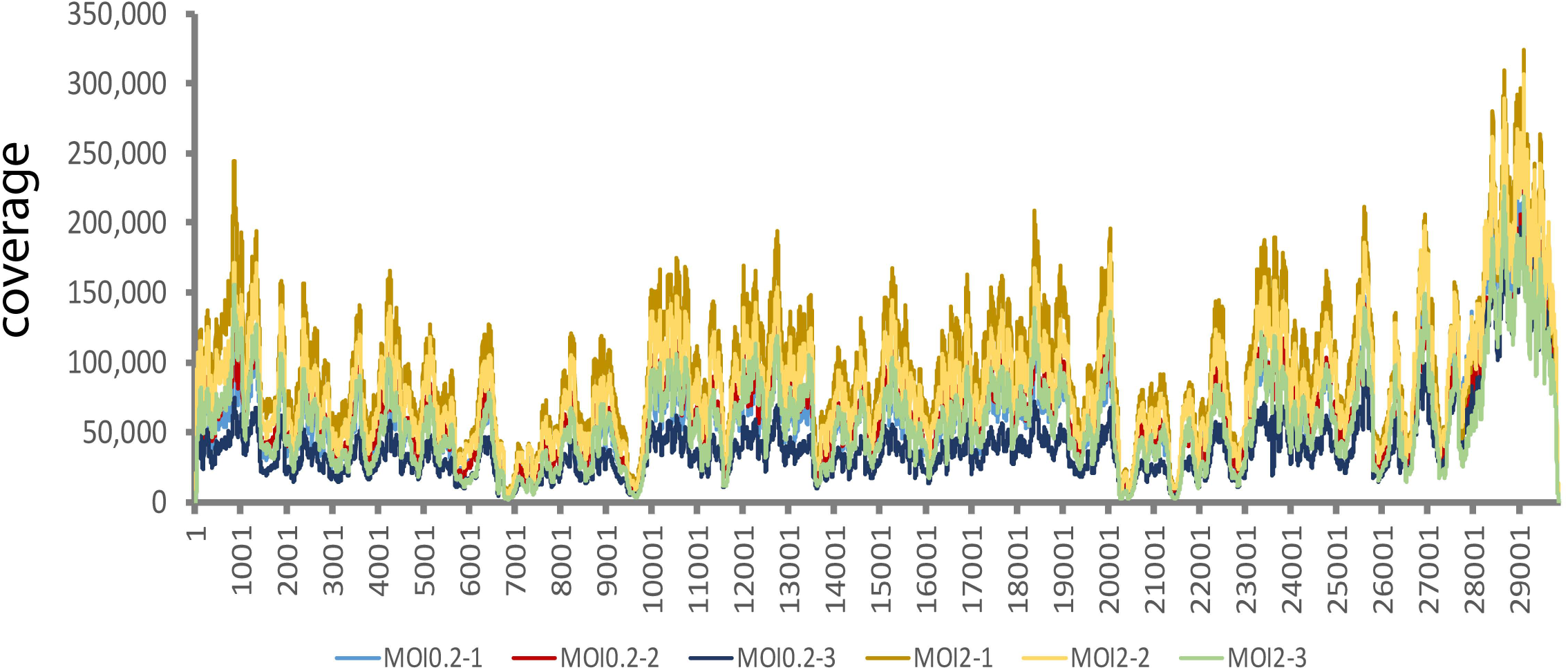
Genome coverage of SARS-CoV-2 infected HAE cells with MOIs of 0.2 and 2, respectively. Six total RNA samples, as indicated with six colors, extracted from HAE-ALI^B2-20^ cultures infected with SARS-CoV-2 with at MOIs of 0.2 and 2, respectively, were subjected to whole RNA-seq. The reads were mapped to the reference SARS-CoV-2 Wuhan-Hu-1 strain genome (MN908947, NCBI), as shown with nucleotide numbers (X axis), using BWA and the sequencing read coverage (Y axis) was calculated.

We further analyzed the viral sgRNA expression in infected HAE cells. Junction-spanning reads covering the 5’-leader and different sgRNAs were counted and analyzed (**Supplemental Material S1**). Different sgRNAs were abundantly expressed in infected HAE cells. As the negative-strand intermediates account only ∼1 % as abundant as their positive sense counterparts (22,37), indicating most of the identified sgRNAs were +sgRNAs. N protein encoding RNA was the most abundantly expressed viral transcript and accounted for 23.11% and 16.93% of total junction-spanning reads in the groups of MOI 0.2 and MOI 2, respectively, followed by ORF3a, ORF7a, M, ORF8, S, E, ORF6 coding RNAs (**Fig. 3**). The junction-spanning reads associated with ORF7b and ORF9a/b were identified at a level of 0.01% or less of the total junction-spanning reads and were only identified in part of all the six samples in two MOI groups (**Supplemental Material S1)**. In SARS-CoV-2 infected HAE cells, S RNA transcript was expressed at a ratio of ∼2% of total junction-spanning reads in both groups (**Fig. 3**), compared to that of ∼8% in Vero cells. We detected relatively higher level of ORF3a (∼8%) transcript in SARS-CoV-2 infected HAE cells (**Fig. 3**), in contrast to 5.22% in infected Vero cells (15).

**Fig. 3.**
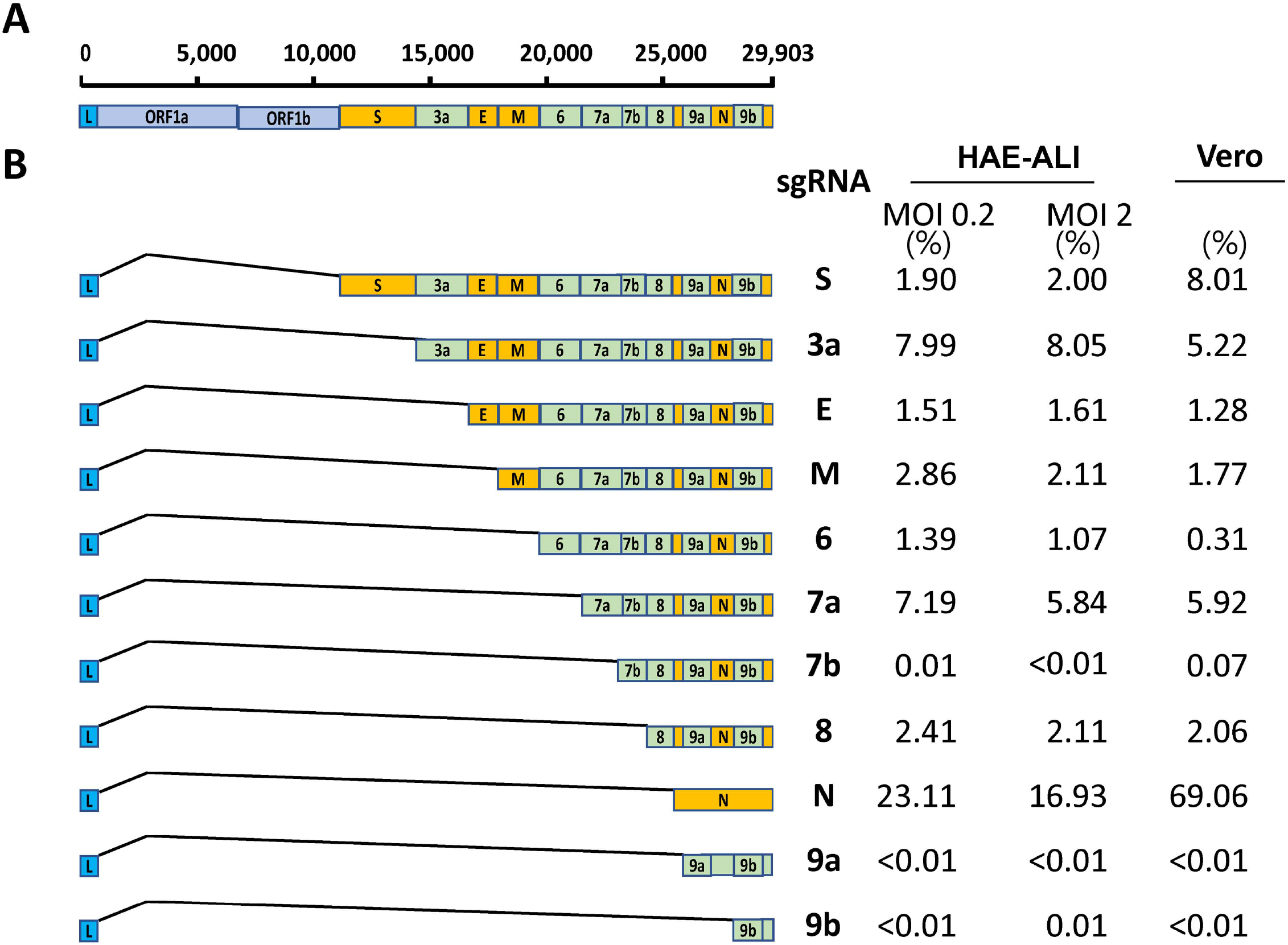
Identification and quantification of SARS-CoV-2 subgenomic RNAs. **(A) Genome organization.** The SARS-CoV-2 genome is schematically diagrammed (not to scale) with regions in order coding for open reading frame 1a (ORF1a)/ORF1b, S protein, ORF3a, E and M proteins, ORF7a/b and ORF8, N protein, and ORF9a/b. The leader sequence was labeled as L in blue box. The structural genes are labeled within boxes in orange and the accessory genes are labeled within boxes in light green. **(B) Subgenomic RNAs**. Six total RNA samples were extracted from SARS-CoV-2 infected HAE-ALI cultures (at MOIs of 0.2 and 2, respectively) and subjected to whole RNA-seq. Three repeats in each MOI group were merged. Junction-spanning reads were identified using STAR (2.7.3a), and the transcript abundance, as shown in % under HAE-ALI/MOI 0.2 or 2, was estimated by counting the reads that span the junction of the corresponding RNA transcript. The left is the diagrammed subgenomic RNAs. The canonical junction-spanning reads related to each sgRNA were calculated and the ratios are shown on right. The abundances of the subgenomic transcripts identified in Vero cells in a previous study (15) are listed for comparison.

Interestingly, in all identified spanning-junction reads, only ∼50% correlated to the canonical sgRNA transcripts in both MOI infection groups. The other half junction-spanning reads represent either reads covering 5’-leader sequence but with unexpected 3’ sites located in the middle of annotated ORFs or reads covering between different ORFs or inside an ORF without 5’-leader sequence (**Supplemental Material S1**). It’s important to note that a lot of these noncanonical junction-spanning patterns were supported by only one read from the RNA-seq data, indicating that these noncanonical transcripts may arise from erroneous replicase activity.

### Identification of deletions surrounding the furin cleavage site at S1/S2

Among the ∼50% noncanonical junction-spanning reads, we identified a high abundant 36-bp deletion, mut-del1, located at nt 23,594-23,629 spanning the FCS (**Fig. 4A**) that encodes aa ^678^TNSP<u>RRAR↓S</u>VAS^689^ (**Fig. 4B**, “↓” indicates cleavage). It displayed at frequencies of 21.04% and 14.79% of total junction-spanning reads in MOI 0.2 and MOI 2 groups, respectively (**Tab. 2**). Another 15-bp deletion, mut-del2, located at nt 23,583-23,597, encoding aa^675^QTQTN^679^ (**Fig. 4A**), just two amino acids ahead of the FCS. It accounted for 0.42% and 15.11% of the total junction-spanning reads in MOI 0.2 and MOI 2 groups, respectively (**Tab. 2**). The ratio of mut-del1 is only slightly lower than the N sgRNA and nearly 10 times higher than the S sgRNA transcript, indicating a high fraction of this mutation comes from the viral genome (+gRNA).

**Fig. 4.**
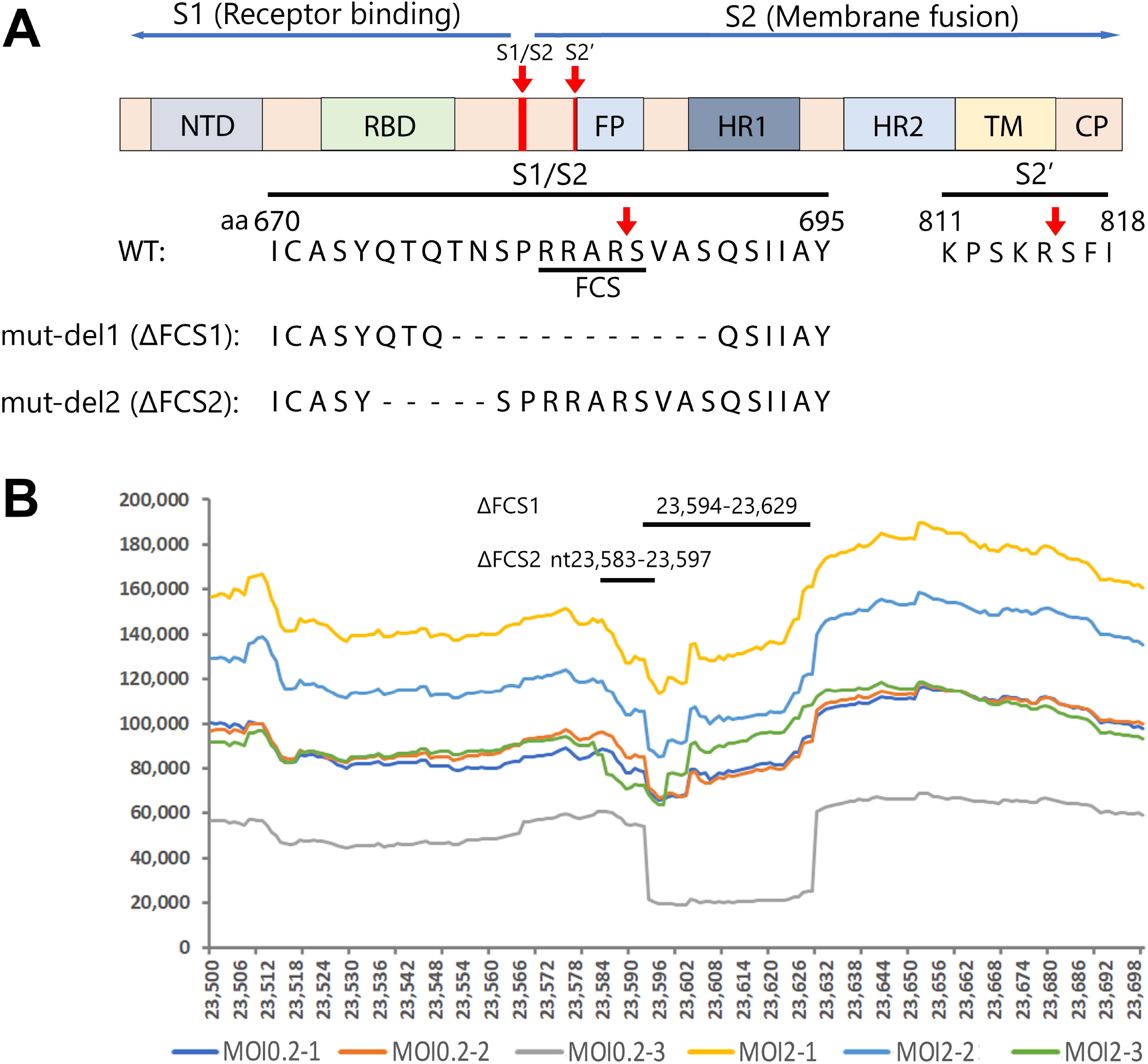
Features of the S gene of SARS-CoV-2 and the deletions detected in the FCS region. **(A) S gene and FCS.** Key domains of the S polypeptide are diagrammed in the context of the SARS-CoV-2 genome. The S1 protein, receptor binding unit, harbors N-terminal domain (NTD) and receptor binding domain (RBD) subunit, which is conserved and recognizes ACE2. The S2, membrane fusion subunit, has fusion peptide (FP), S2’ proteolytic site, two heptad-repeats, HR1 and HR2, and a transmembrane domain (TM) followed by cytoplasmic peptide (CP) (30). The S protein has acquired a polybasic site (RRAR↓S, a furin cleavage site, FCS) for cleavage at S1/S2 boundary. An FCS region of aa670-695, together with the two key deletions mut-del1 (ΔFCS1) and mut-del2 (ΔFCS2), are shown with S amino acid sequences of the SARS-CoV-2 genome (GenBank, accession code MN908947). **(B) Coverage plots of S gene at nt 23**,**500 to 23**,**698 in SARS-CoV-2 infected HAE-ALI**^**B2-20**^. The coverage plots show the most abundant junction-spanning reads in SARS-CoV-2 infected HAE-ALI^B2-20^ cultures are the 36 bp and 15 bp deletions in S gene of nt 23,594-23,629 and nt 23,583-23,597, respectively, which deleted 12 aa and 5 aa shown in mut-del1 and mut-del2 in panel A. RNA Sample 5, 6, and 7 were extracted from HAE-ALI^B2-20^ infected at an MOI of 0.2 at 4 dpi, and RNA Sample 9, 10, and 11 were extracted at MOI of 2 at 4 dpi.

**Table 2.** Ratio of reads covering mut-del1 and mut-del2 to total junction spanning reads and viral genome.

|  | Junction-spanning reads (%)<sup>a</sup> |  | Viral genome ratio (%)<sup>b</sup> |  |
| --- | --- | --- | --- | --- |
|  | MOI 0.2 | MOI 2 | MOI 0.2 | MOI 2 |
| **Mut-del1** | 21.04 | 14.79 | 69.02 | 20.02 |
| **Mut-del2** | 0.42 | 15.11 | 1.11 | 15.75 |

Note: a, the minimal size of the junctions was set 10 as described in Materials and Methods. b, the total reads include both viral genome RNA (gRNA) and subgenomic RNA (sgRNA).

**Table 3.** Summary of the detections of mut-del1 and mut-del2 in stock viruses and apical washes of SARS-CoV-2 infected HAE-ALI cultures derived from different donors.

| Virus or donor | dpi (source) | Mut-del1 |  | Mut-del2 |  |
| --- | --- | --- | --- | --- | --- |
|  |  | RNA-seq (%) | PCR-seq (%) | RNA-seq (%) | PCR-seq (%) |
| **P0 stock** | (BEI) | 0.87 | 2.03 | 2.09 | 2.45 |
| **P1 stock** | (Vero-E6) | 21.69 | 40.47 | 5.18 | 5.16 |
| **B3-20** | 3 | ND |  | 23.17 |  |
|  | 12 | ND |  | 4.44 |  |
|  | 20 |  | 0.00 |  | 0.57 |
| **B4-20** | 3 | 1.08 |  | 8.33 |  |
|  | 12 |  | 0.052 |  | 4.27 |
|  | 13 |  | 0.033 |  | 3.90 |
|  | 17 |  | 0.103 |  | 1.59 |
| **B9-20** | 5 |  | 0.001 |  | 0.01 |
|  | 11 |  | 0.064 |  | 0.09 |
|  | 17<sup>#</sup> |  | 0.056 |  | 0.15 |
|  | 17<sup>#</sup> |  | 0.024 |  | 0.07 |
| **KC19** | 4 | 0.16 | 1.79 | 20.75 | 35.71 |
|  | 13 | ND |  | 20.98 |  |
|  | 21 |  | 0.017 |  | 41.79 |
| **L209** | 3 |  | 0.037 |  | 30.33 |
|  | 14 |  | 0.001 |  | 0.11 |
|  | 41 |  | 0.001 |  | 0.04 |

ND, not detected or <0.1%. # independent samples.

To further reveal the ratio of the two deletions in total viral genome, the junction-spanning reads associated with the two deletions were normalized with the average reads covering the same deletions. The results showed that 69.02% and 20.02% of the viral reads related to this region contain the mut-del1 deletion, while 1.11% and 15.75% of this region contain mut-del2 deletion in MOI 0.2 and MOI 2 groups, respectively (**Tab. 2**). It should be noted that the total reads used for normalization include reads of both viral gRNA and sgRNAs. Thus, here we were unable to distinguish the origin of these two deletions from the viral genome and viral RNA transcripts in these total cellular transcriptome data.

Except for these two highly abundant mut-del1 and mut-del2 deletions, we also observed a 21-bp FCS deletion at nt 23,595-23,615, encoding aa ^678^TNSPRRA^684^, but only in the MOI 2 group with 1.13% of the total junction-spanning reads, and a 39-bp deletion at the N-terminus of the S protein (nt 21,743-21,781 encoding aa ^61^NVTWFHAIHVSGT^73^) with 0.27% and 0.60% of the total junction-spanning reads in MOI 0.2 and MOI 2 groups, respectively.

In addition to these deletions in S gene, we identified about 50 different in-frame or frameshift deletions in M encoding region that appeared in all six samples of both MOI groups, and there were even more deletions in M coding region that appeared in only a part of the six RNA samples (**Supplemental Material S1**). Although the ratio of single deletion was low, the 50 deletion patterns that appeared in all 6 RNA samples had the ratios of 2.39% and 3.18% in MOI and MOI 2 groups, respectively, which is similar or even higher than the identified canonical junction-spanning reads related to M sgRNAs (**Fig. 3**). Notably, most of these identified deletion patterns of M gene also appeared in SARS-CoV-2 infected Vero cells (15). Whether these deletions produce functional M protein or affect the function of M protein warrant further studies. In SARS-CoV-2 infected Vero-E6 cells, a high ratio of 27-bp deletion in E gene (nt 26,257-26,283) was identified (15), which, however, was not found in infected HAE cells.

### Dynamics of the FCS region deletions in virions apically released from SARS-CoV-2 infected HAE-ALI cultures derived from various donors

To further investigate the FCS region deletions during SARS-CoV-2 infection of HAE cells, we infected HAE-ALI cultures generated from five different donors, B3-20 (MOI=0.2), B4-20 (MOI=2), B9-20 (MOI=2), L209 (MOI=0.2) and KC19 (MOI=0.2), and collected the progeny in the apical washes at different time points. The dynamics of apical virus releases of the HAE-ALI cultures of B3-20, B4-20, B9-20, and L209 have been described in our previous study (36). The apical virus release kinetics of the HAE-ALI^KC19^ is shown in **Fig. 5**. Viral RNA was prepared either for RNA-seq or for PCR amplicon-seq of a 384-nt sequence covering the FCS. Notably, mut-del1 was not significantly detected (<0.1%) in all the apically released viruses collected at >13 dpi (**Tab. 3**, Bx-20). Nevertheless, for viruses collected from HAE-ALI^KC19^, the mut-del2 was detected at a high level (20.75%^RNA-seq^ and 20.98%^RNA-seq^) at 4 dpi and 13 dpi, respectively, which reached a close level of 41.79%^PCR-seq^ at 21 dpi. Although the viruses derived from HAE-ALI^B3-20^, HAE-ALI^L209^ and HAE-ALI^B4-20^ contain a high level (23.17%, 30.33%, and 8.3%, respectively) of mut-del2 at 3 dpi, it decreased to a level of <2% at ≥17 dpi (**Tab. 3**, Bx-20). HAE-ALI^B9-20^ did never produce significant mut-del2 (<0.1%).

**Fig. 5.**
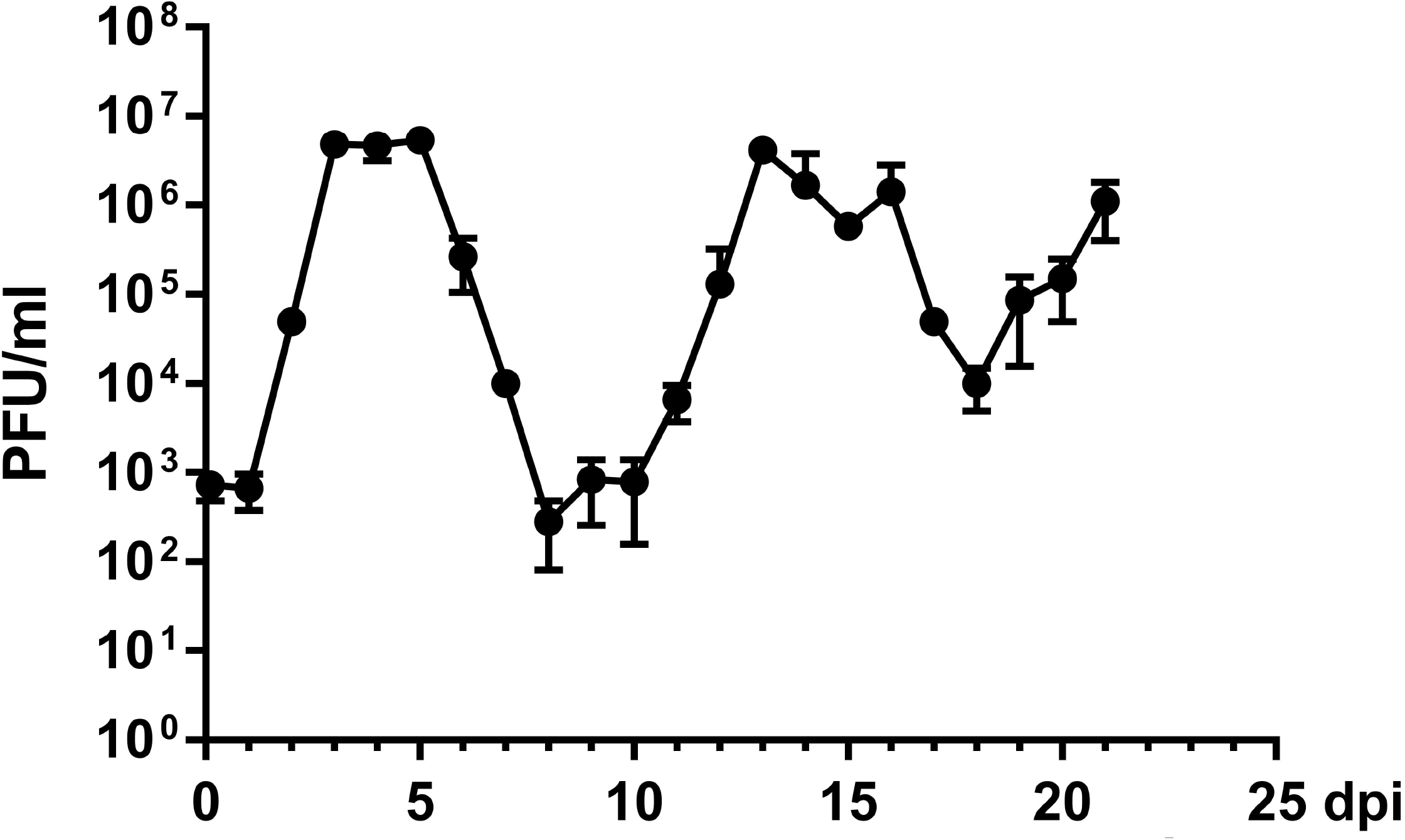
Apical virus release kinetics of SARS-CoV-2 infected HAE-ALI^KC19^ culture. HAE-ALI^KC19^ cultures were infected with SARS-CoV-2 at an MOI of 0.2 from the apical side. At the indicated days post-infection (dpi), the apical surface was washed with 300 µl of D-PBS to collect the released viruses. Plaque-forming units (PFU) were determined (y axis) and plotted to the dpi. Values represent means ± standard deviations (SD) (error bars).

Notably, SARS-CoV-2 isolate USA-WA1/2020 P0 stock provided by BEI was already passaged 4 times in Vero cells, and it was reported that there appeared significant heterogeneity at the S1/S2 boundary in Vero cell propagated virus (31). To verify this, we sequenced the viral RNA of P0 (the originally received vial) and P1 (passaged once in Vero-E6 cells) virus stocks. The results showed that there was no detectable mut-del1 in the P0 stock but a high rate of 21.69%^RNA-seq^ and 40.47^PCR-seq^ in the P1 stock. However, while there was ∼2% of mut-del2 in the P0 stock, it only slightly increased to ∼5% in the P1 stock. (**Tab. 3**, P0 and P1). These results confirmed that even though there was no or low level of mut-del1 and mut-del2 in the P0 stock, there was a high level of mut-del1 and a low level of mut-del2 in the P1 stock, which we used for infection of all HAE-ALI cultures.

Taken together, the results demonstrated that the mut-del1 appears at a very low rate in all the HAE-ALI produced viruses at late time points of infection (≥17 dpi), which was confirmed by both RNA-seq and PCR amplicon-seq. Although the inoculum had the mut-del1 detected at a high frequent rate of 21.69%^RNA-seq^ (40.47%^PCR-seq^), this deletion was obviously suppressed when the virus propagated in the HAE-ALI cultures. In addition, except for the viruses produced from HAE-ALI^KC19^, mut-del2 also appeared at a low detection rate during the course of infection (≥17 dpi) in various HAE-ALI cultures.

### FCS region deletions during SARS-CoV-2 infection of human airway epithelia are donor dependent

The above RNA-seq and PCR-seq results of apically released virions from five individual HAE-ALI cultures suggested a selective pressure in suppressing the deletions of the FCS region during SARS-CoV-2 propagation in human airway epithelia. The exception is the infection in HAE-ALI^KC19^ cultures, which amplified the mut-del2 to a high level. To address the possibility of the donor dependency of the FCS region deletions, we infected HAE-ALI cultures generated from two additional donors B15-20 and B16-20, and collected both the viral progeny in apical washes at the early and late time points post-infection for PCR-seq. The apical washes from infected HAE-ALI^B15-20^ and HAE-ALI^B16-20^ had virus titers of > 10^6^ pfu/ml at both 3 dpi and 13 dpi, indicating the input inoculum replicated similarly in these two HAE-ALI cultures as in the others (**Fig. 5**) (36). The sequencing results of the apically released viruses from infected HAE-ALI^B15-20^ showed that mut-del1 was detected at a rate of 31.87% at 3 dpi, which increased to a high level at 54.22% at 13 dpi (**Tab. 4**, B15-20/mut-del1). However, mut-del2 was barely detectable at a low rate (0.18%) at 3 dpi, and this rate remained very low at 0.69% at 13 dpi (**Tab. 4**, B15-20/mut-del2). For the viruses apically released from infected HAE-ALI^B16-20^ cultures, only mut-del1 was detected at a rate of 6.81% at 3 dpi, which nearly disappeared (at rate of 0.08%) at 13 dpi; whereas mut-del2 was detected at very low levels (<1%) at both 3 and 13 dpi (**Tab. 4**, B16-20). The suppression of mut-del1 in HAE-ALI^B16-20^ cultures was similar as what observed in the previously tested five ALI cultures (**Tabl. 3**).

**Table 4.** Detections of mut-del1 and mut-del2 in SARS-CoV-2 virions apically released from infected HAE-ALI cultures derived from B15-20 and B16-20 donors using PCR-seq.

| Donor | dpi | Mut-del 1 | Mut-del2 |
| --- | --- | --- | --- |
|  |  | PCR-seq | PCR-seq |
| **B15-20** | 3 | **31.87** | 0.18 |
|  | 13 | **54.22** | 0.69 |
| **B16-20** | 3 | 6.78 | 0.71 |
|  | 13 | 0.08 | 0.16 |

Together with the detection rates of the viruses produced from the infected HAE^KC19^-ALI cultures, the above results suggested that the probability of FCS region deletions is dependent on the HAE-ALI cultures made from airway epithelial cells of different donors. In contrast to the HAE-ALI^KC19^ that produced a high rate of mut-del2, HAE-ALI^B15-20^ tended to generate the viruses that have a high rate of the mut-del1 deletion.

## Discussion

In this study, we analyzed the transcriptome of SARS-CoV-2 in polarized human bronchial airway epithelia, an in vitro model mimicking the SARS-CoV-2 infection in human lower airways (35,36). We found that the transcriptome in HAE-ALI reflects more closely the viral transcriptome in the airways of COVID-19 patients, supporting that HAE-ALI is a physiologically relevant in vitro culture to study SARS-CoV-2. Neither RNA-seq data of clinical SARS-CoV-2 positive nasopharyngeal specimens nor RNA-seq of SARS-CoV-2 infected HAE-ALI showed the 5’-leader sequence read peak (38,39). In SARS-CoV-2 infected Vero-E6 cells, N sgRNAs accounted for up to 69% in total viral RNA transcripts, and S sgRNAs accounted for 8% of the total junction-spanning reads (15) (**Fig. 3**). Nevertheless, in HAE-ALI cultures, SARS-CoV-2 still expresses the abundant N protein transcript, and relatively low level of S gene mRNA. Of note, the overall sgRNA transcripts in infected HAE-ALI were mapped to only 50% of all the canonical sgRNAs, much lower than that in Vero cells (15), which is partially due to the high deletion rate of the FCS region derived from the inoculated virus (P1 stock, **Tab. 3**).

Among all the SARS-CoV-2 viral genes, S gene is the one most variable, in particular the S1/S2 junctional region, featuring the FCS. Increasing evidence has shown that the S1/S2 FCS region is highly unstable, and various deletions and mutations have been detected or isolated in SARS-CoV-2 infected Vero cells (17,31-34). A mutant with a 30-bp deletion, encoding aa ^679^NSPRRAR↓SVA^688^, showed enhanced replication ability in Vero cells, and had the capability to dominate the genome population during passage in Vero cells (34). A 21-bp deletion encoding aa ^679^NSPRRAR^686^ were detected (>10%) in low (<2-3) passaged isolates (32). However, detection of the original clinical specimen where the mutant was derived and SARS-CoV-2 positive clinical specimens showed no such deletions (15-30 bp) in the FCS region (33), indicating that the mut-del1 or mut-del1-like (containing FCS) deletions are generated during the propagation in Vero cells. Apparently, SARS-CoV-2 is under strong selection pressure in Vero cells to acquire adaptive mutations in the S protein. Nevertheless, mut-del2 (^675^QTQTN^679^) has not only been identified in Vero cell passaged isolates (40), but also in 3 of 68 clinical specimens (32), indicating that mut-del2 maybe clinically more important (relevant) than mut-del1.

The S protein as a part of the viral envelope facilitates viral entry into infected cells. The S1 subunit contains the receptor binding domain and the S2 domain mediates fusion of the viral envelope with a cellular membrane (23). The infectivity of SARS-CoV-2 necessitates the activation of S protein. There are two proteolytic activation events associated with S-mediated receptor binding and membrane fusion. The first is a priming cleavage that occurs at the S1/S2 boundary, and the second is the obligatory triggering cleavage that occurs within the S2’ site (**Fig. 4A**). The priming cleavage at S1/S2 boundary causes the conformation changes of the S1 subunit for receptor binding and of the S2 subunit for conversion of a fusion competent form, by enabling the S protein to better bind receptors or expose the hidden S2’ cleavage site. The cleavage at S2’ triggers the fusion of the viral envelope with the host cell membrane (23). Cleavage by furin at the S1/S2 site is required for subsequent transmembrane serine protease 2 (TMPRSS2)-mediated cleavage at the S2’ site during viral entry into lung cells (41). However, a cathepsin B/L-dependent auxiliary activation pathway is available during infection of SARS-CoV-2 infection in TMPRSS2 negative cells (34,42), which is likely not dependent on the cleavage at S1/S2 (43). One important novel finding of our study is that HAE-ALI cultures prepared from human airway cells isolated from different donors selected different FCS deletions. While most (5/7) of the HAE-cultures (B3-20, B9-20, L209, and B16-20) strongly selected the FCS during virus replication (the FCS deletions only accounted <1% at 13 dpi), HAE-ALI^KC19^ prefers selection of mut-del2 (^675^QTQTN^679^) (41.79%^PCR-seq^ at 21 dpi), and HAE-ALI^B15-20^ selected the mut-del1 (^678^TNSP<u>RRAR↓S</u>VAS^689^) at a rate of 54.22% ^PCR-seq^ at 13 dpi. Although mut-del2 retains the FCS, deletion of QTQTN upstream of the FCS also prevented the cleavage (40). These mutants with amino acid deletions immediately upstream of FCS, like mut-del2, or downstream (^685^RSV^687^ or ^689^SQS^691^) also showed significant defects in S protein processing (40,42). Both types of the FCS region deletions were unable to utilize the furin and TMPRSS2-mediated plasma membrane fusion entry pathway and exhibited a more limited range of cell tropism (42,44). This is substantiated by the fact that there were no FCS region deletions detected in SARS-CoV-2 propagated in TMPRSS2-expressing cells (42). Overall, we believe that human airway epithelial cells express ACE2 and TMPRSS2 (36,45-47), which plays an important role during S protein priming and viral entry, and the virus entry is mediated by the membrane fusion pathway. However, from two out of the seven HAE-ALI cultures tested in this study, the lack of suppression of the FCS region deletion was also found in apically released virions of the infected HAE-ALI cultures made from KC19 and B15-20 donors. Previously, we discovered that the SARS-CoV-2 infection in HAE-ALI resulted in periodic recurrent replication peaks of progeny (36). Since the cleavage at the S2’ site by TMPRSS2 necessitates the priming cleavage at S1/S2, the accumulation of FCS mutations in the progeny during the infection in HAE-ALI^B15-20^ and HAE-ALI^KC19^ suggests they are more permissive to the infection of the FCS-deficient SARS-CoV-2 mutants than the other cultures. We speculate that epithelial cells from these two donors may express much less TMPRSS2, and therefore the virus utilizes the TMPRSS2-independent and cathepsin-dependent endosomal entry pathway (42,44,48), which likely does not require the S cleavage at S1/S2 (43) and thus prefers replication of the FCS region deletion mutants.

Importantly, the deletion of QTQTN (mut-del2) diminished SARS-CoV-2 entry and infection in Vero-E6 cells (40). Furthermore, three FCS-related deletion mutants, ΔPRRA↓, ΔRAR↓SVAS, and ΔNSPRRAR↓SVA, have been shown to have reduced replication in vitro and lung disease in animal models (44,49,50), strongly supporting that the FCS is a virulence-related motif. Since the ΔQTQTN also abolished the furin cleavage (40), we speculate mut-del2 mutant should have reduced lung disease in animals as well. Since the FCS is a key motif related to virulence, it is important to investigate the natural occurrence rate of the FCS region deletions, possibility or limitation of their human-to-human transmission, as well as their pathogenicity. Several studies tried to screen the FCS region deletions from patients-derived SARS-CoV-2. As discussed above, screening of 27 SARS-CoV-2 positive clinical specimens, including one specimen that had FCS deletions identified after passaging in Vero-E6 cells, failed to detect any FCS deletions (33). However, one study detected the ^675^QTQTN^679^-deleted mutants (mut-del2) in 3 of 68 SARS-CoV-2 positive clinical specimens (32). In another detection of 51 SARS-CoV-2 positive patient specimens, although a high rate of 52.9% and 82.4% of the positive clinical samples contained the FCS upstream motif (^661^ECDIPIGAG^669^) and the PRRA deletions, respectively, the mutant population is at a very low level (0.33% ±1.17% for FCS upstream motif deletion and 1.12% ±1.21% for PRRA deletion) (51), arguing the infectivity and transmissibility of these mutants.

Along with the usages of antibody drugs and the wide inoculation of the vaccine, which target the S protein, the virus may undergo further mutations under the pressure of human immune response. Supervision and screening the mutations in the S protein gene in clinical specimens is extremely important to identify the escaped isolates which may increase or decrease infectivity and transmissibility. Apparently, the in vitro polarized HAE model, which can facilitate long-term infection of SARS-CoV-2, is an ideal model to study S gene mutants under various conditions.

## Acknowledgments

We thank the Cells and Tissue Core of Center for Gene Therapy, the University of Iowa (DK054759) for providing the primary cell cultures. The following reagent was deposited by the Centers for Disease Control and Prevention and obtained through BEI Resources, NIAID, NIH: SARS-Related Coronavirus 2, Isolate USA-WA1/2020, NR-52281. The study was supported by PHS grant AI150877 and an internal award from University of Kansas Medical Center.

